# A global perspective of codon usage

**DOI:** 10.1101/076679

**Authors:** Bohdan B. Khomtchouk, Claes Wahlestedt, Wolfgang Nonner

## Abstract

Codon usage in 2730 genomes is analyzed for evolutionary patterns in the usage of synonymous codons and amino acids across prokaryotic and eukaryotic taxa. We group genomes together that have similar amounts of intra-genomic bias in their codon usage, and then compare how usage of particular different codons is diversified across each genome group, and how that usage varies from group to group. Inter-genomic diversity of codon usage increases with intra-genomic usage bias, following a universal pattern. The frequencies of the different codons vary in robust mutual correlation, and the implied synonymous codon and amino acid usages drift together. This kind of correlation indicates that the variation of codon usage across organisms is chiefly a consequence of lateral DNA transfer among diverse organisms. The group of genomes with the greatest intra-genomic bias comprises two distinct subgroups, with each one restricting its codon usage to essentially one unique half of the genetic code table. These organisms include eubacteria and archaea thought to be closest to the hypothesized last universal common ancestor (LUCA). Their codon usages imply genetic diversity near the hypothesized base of the tree of life. There is a continuous evolutionary progression across taxa from the two extremely diversified usages toward balanced usage of different codons (as approached, e.g. in mammals). In that progression, codon frequency variations are correlated as expected from a blending of the two extreme codon usages seen in prokaryotes.

**AUTHOR SUMMARY:** The redundancy intrinsic to the genetic code allows different amino acids to be encoded by up to six synonymous codons. Genomes of different organisms prefer different synonymous codons, a phenomenon known as ‘codon usage bias.’ The phenomenon of codon usage bias is of fundamental interest for evolutionary biology, and is important in a variety of applied settings (e.g., transgene expression). The spectrum of codon usage biases seen in current organisms is commonly thought to have arisen by the combined actions of mutations and selective pressures. This view focuses on codon usage in specific genomes and the consequences of that usage for protein expression.

Here we investigate an unresolved question of molecular genetics: are there global rules governing the usage of synonymous codons made by genomic DNA across organisms? To answer this question, we employed a data-driven approach to surveying 2730 species from all kingdoms of the ‘tree of life’ in order to classify their codon usage. A first major result was that the large majority of these organisms use codons rather uniformly on the genome-wide scale, without giving preference to particular codons among possible synonymous alternatives. A second major result was that two compartments of codon usage seem to co-exist and to be expressed in different proportions by different organisms. As such, we investigate how individual different codons are used in different organisms from all taxa. Whereas codon usage is generally believed to be the evolutionary result of both mutations and natural selection, our results suggest a different perspective: the usage of different codons (and amino acids) by different organisms follows a superposition of two distinct patterns of usage. One distinction locates to the third base pair of all different codons, which in one pattern is U or A, and in the other pattern is G or C. This result has two major implications: (1) the variation of codon usage as seen across different organisms is best accounted for by lateral gene transfer among diverse organisms; (2) the organisms that are by protein homology grouped near the base of the ‘tree of life’ comprise two genetically distinct lineages.

We find that, over evolutionary time, codon usages have converged from two distinct, non-overlapping usages (e.g., as evident in bacteria and archaea) to a near-uniform, balanced usage of synonymous codons (e.g., in mammals). This shows that the variations of codon (and amino acid) biases reveal a distinct evolutionary progression. We also find that codon usage in bacteria and archaea is most diverse between organisms thought to be closest to the hypothesized last universal common ancestor (LUCA). The dichotomy in codon (and amino acid usages) present near the origin of the current ‘tree of life’ might provide information about the evolutionary development of the genetic code.

## INTRODUCTION

The standard genetic code associates 18 of the 20 different amino acids redundantly with two to six synonymous codons. Usage of different synonyms thus can in principle encrypt information beyond that of protein primary structure [1]. Indeed, different codons specifying the same amino acid occur with diverse frequencies within genes, within genomes, and across genomes.Cryptic information contained in the DNA of genes (and the transcribed mRNA) could in a variety of ways direct mRNA processing and/or translation [1], possibilities that have motivated investigations of the evolution and consequences of ‘codon usage bias’ (reviewed by [2–4]).

Molecular restoration studies of extinct forms of life, as originally proposed by Pauling and Zuckerkandl [5] and now pursued in the reconstruction of phylogenetic trees, are based on the primary structure information encoded in DNA [6, 7]. Forms of life close to the base of the tree (the hypothetical last universal common cncestor (LUCA)) have been inferred from homologous proteins found in free-living bacteria and archaea [8, 9]. On the other hand, the origin of the genetic code that links genome and proteome is still elusive, much as it was when the code was discovered [10, 11].

The study described in this paper originated from the observation that genomes of certain bacteria reveal two nearly exclusive usages of the genetic code. A profound genetic divergence near the putative base of the tree of life is potentially important for inferences on the evolutions of codon usage and, ultimately, of the genetic code. We therefore survey intragenomic bias and between-genome diversity of codon usage over different taxa using the codon-usage compilation of the Codon Usage Tabulated from GenBank (CUTG) database [12]. Our results indicate that the variation of codon usage across genomes is chiefly a variation that is correlated over most different codons, swaps the nucleotide in the third position of codons (U⇄C, or A⇄G), and alters amino acid usage, as well. Both bias and diversity of codon usage decrease from prokaryotes to multicellular organisms. Whereas the majority of the sampled genomes have small bias and diversity of codon usage, genomes of prokaryotes thought to be close to the base of the tree of life are highly biased and form two distinct genetic lineages.

## METHODS

We analyze genomes that are represented in the CUTG database with codon counts 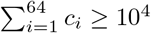 (database files ‘gbxxx.spsum.txt’ where xxx = bct, inv, mam, phg, pln, pri, rod, vrl, vrt). Entries for mitochondrial or other organelle genomes are excluded. The count *c_i_* of occurrences of a different codon *i* (1 ≤ *i* ≤ 64) in an analyzed genome is converted to the frequency

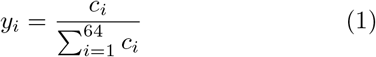

On the basis of frequencies we compute non-parametric measures to quantify variations of codon usage within and between genomes. To describe codon usage within a particular genome, we take the frequencies of the 64 different codons as the coordinates of that genome in a 64-dimensional Euclidean space. The Euclidean distance between that point and a reference point representing a genome using the 64 different codons with equal frequencies then defines a measure of frequency variation within the genome, which we call ‘bias’ *B*:

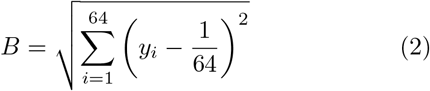

For a hypothetical genome in which all different codons occur with equal frequencies, bias *B* = 0. Real genomes will have bias *B* > 0: even though synonymous codons encoding for the same amino acid might be used equally, the amino acid stoichiometry of an encoded proteome requires non-uniformity in the usage of different codons.

We measure the variation of usage of a different codon *i* in a set of genomes *k* = 1,…,*N* by the variance

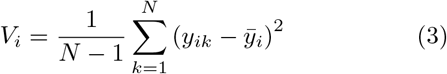

where 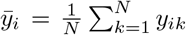 is the average frequency of the different codon *i* in the set of *N* genomes. As a global measure of how usage of all different codons varies in the set of genomes, we define ‘diversity’ of codon usage as the total variance

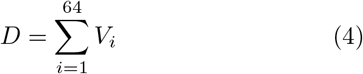

Diversity in a set of genomes will be zero if all genomes have zero bias. Diversity will also be zero if bias is not zero but all genomes *k* of the set use particular different codons *i* with the same frequencies, *y_ik_* = *y_i_*.

In order to detect dependencies among different codons in the variation of codon usage, we describe diversity more specifically by the 64 × 64 covariance matrix with the elements

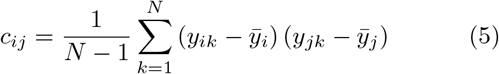

The covariance matrix is subjected to singular value decomposition (SVD) to determine its eigenvalues and eigenvectors [13]. The eigenvectors of the covariance matrix are orthonormal. They define linear combinations of variations in the original codon frequencies that are mutually independent of one another and account for amounts of variance (diversity) that are quantified by the associated eigenvalues. The purpose of the analysis is to find joint variations of codon frequencies (principal components) that make large contributions to the diversity.

As will be described in Results, the eigenvector associated with the largest eigenvalue (first principal component) accounts for much of the observed total diversity of codon usage. We therefore focus further analysis on that first principal component. We compute the projection of the observed codon frequencies *y_i_* of genome k onto the axis defined by the first eigenvector, (*V_i_^1^*):

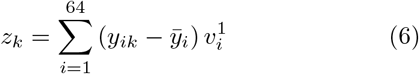

To the extent that the first principal component accounts for the diversity in a set of genomes, the projection *z_k_* describes the codon usage of an entire genome *k* by a single coordinate, which we will use to generate a map of codon usage. The individual codon frequencies of the genome are reconstructed from the first principal component:

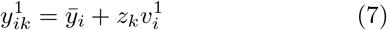

These frequencies are compared with the observed original codon frequencies *y_ik_* to assess the completeness of the description by one principal component.

For a second test of the description by the first principal component, we compute the bias of the codon frequencies reconstructed from the first principal component. Inserting the frequencies 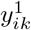 from eq. 7 into eq. 2 yields

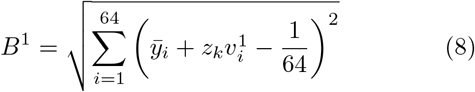

In order to assess variations of amino acid usage that are implied by the variations of codon frequencies we also identify principal components for the amino acid frequencies. Those 20 frequencies and that of the stop signal are computed as partial sums of codon frequencies taken over the groups of codons encoding the different amino acids and the stop signal. In that ‘translation’, we use the standard genetic code, ignoring variants known for some organisms.

## RESULTS

Frequencies of codon occurrence in genomic DNA of *Homo sapiens* and *Arabidopsis thaliana* extracted from the CUTG database (see Methods) are shown in Fig. 1A. The human frequencies have been arranged into a descending sequence (*red color*). The frequencies of the different codons of the plant when arranged to follow the order of the different codons in the human sequence do not form a descending sequence (*blue color*). When the codon frequencies of the plant are by themselves arranged into a descending sequence, the resulting curve is similar to that for the ranked human sequence (*line*). Particular different codons, however, are used with variable preferences in the human and plant genomes, with balance tipping more or less stochastically one way or the other along the 64 different codons.

**FIG. 1.**
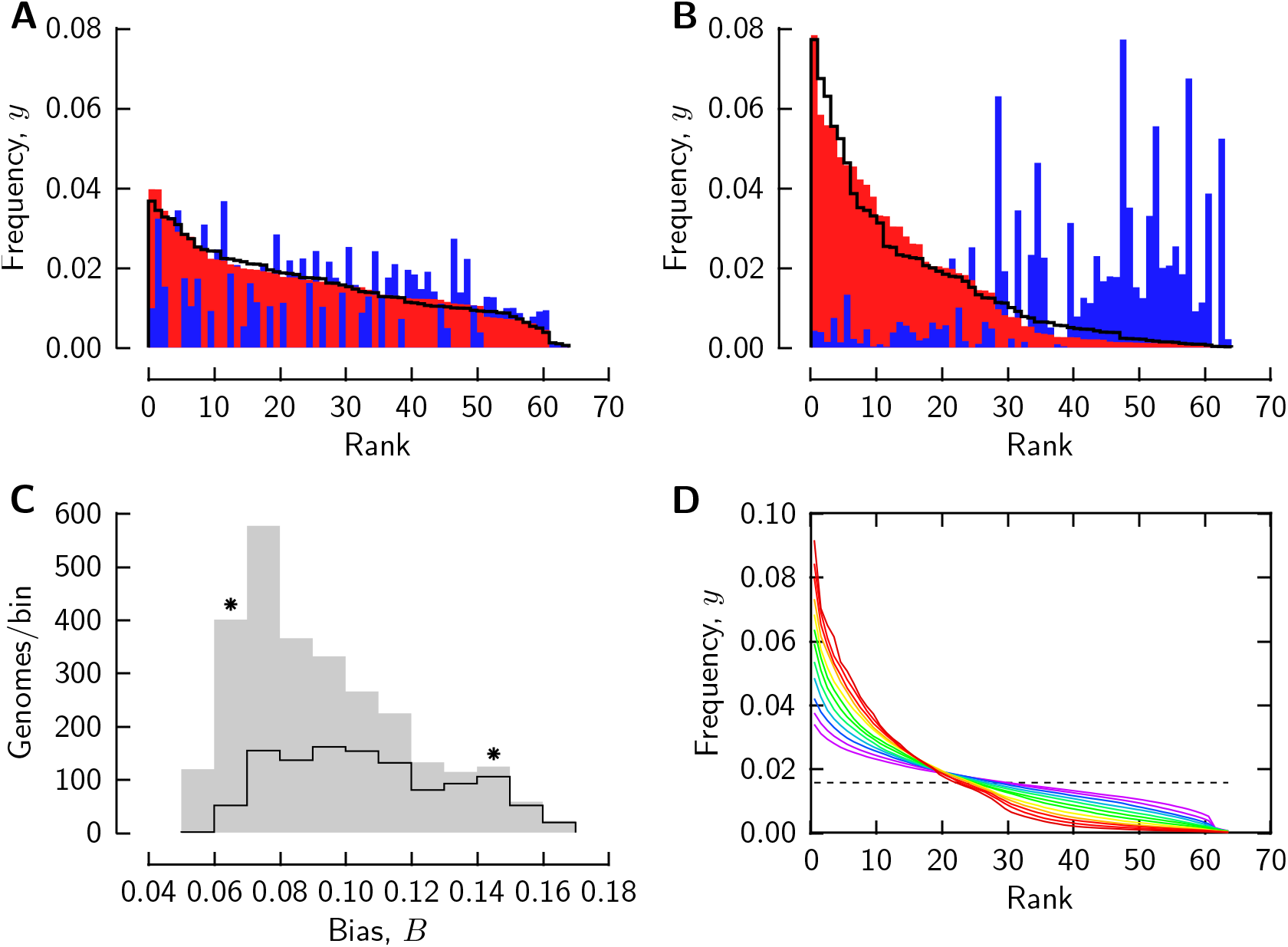
Biased usage of different codons in genomes. Codon frequencies of *Homo sapiens* (A, red color), *Arabidopsis thaliana* (A, blue color), *Streptomyces griseus* (B, red color), and *Clostridium tetani E88* (B, blue color). Frequencies are arranged as described in the text, with the smaller frequency of each two compared genomes shown in the foreground. C: histograms of bias *B* for 2730 genomes from the CUTG database (gray shaded) and of the bacterial genomes in that set (line). Asterisks mark the bins to which the genomes in A,B locate. D: frequency/rank curves averaged over the genomes of each bin in C (with bias increasing from purple to red).

Fig. 1B presents two further sample genomes, of the bacteria *Streptomyces griseus* (*red color*) and *Clostridium tetani E88* (*blue color*), analyzed like the genomes of Fig. 1A. Each bacterial genome uses different codons to much different degrees, but when codon frequencies are individually arranged in descending order, similar frequency/rank curves emerge. On the other hand, the two bacterial genomes encode proteins with nearly exclusive subsets of the 64 different codons.

The four genomes in Fig. 1A,B have been chosen in hindsight of an analysis that is described in the following. They are examples for the variety of codon usages that we found in 2730 genomes from all sections of the CUTG database (see Methods). We will return to the genomes of Fig. 1A,B to illustrate results of that analysis.

We assess non-uniformity of codon usage observed within an individual genome using as a non-parametric measure the ‘bias’ *B* defined by eq. 2 of Methods. Bias measures how much the observed frequencies of a genome diverge from those of a hypothetical genome in which all different codons are used with the same frequency. The histogram in Fig. 1C, summarizing the values of *B* for 2730 genomes of the CUTG database (*gray shade*), peaks near the lower end of the bias range. (The genomes of Fig. 1A,B locate to the bins marked by *asterisks*.) The leftmost four bins comprise 1460 genomes - more than half of the genomes locate to the lower third of the binned range of bias. The majority of genomes use different codons with relatively small bias, like the human and *Arabidopsis* genomes (Fig. 1A), rather than with strong bias, like the bacterial genomes of Fig. 1B. Overall, bacterial genomes use different codons with a wide range of bias, as is apparent when their codon frequencies are binned separately (*line* in Fig. 1C). Although bacteria is the largest CUTG database section, comprising 1134 genomes of the analyzed 2730 genomes, the peak of the overall distribution of bias is mainly due to non-bacterial genomes.

Since frequency/rank curves of genomes have been described or modeled in the past for small sets of genomes ([14–16]), it is of interest to inspect frequency/rank curves from a large genome set as analyzed here. Each curve in Fig. 1D is the average of the frequency/rank curves for the genomes of one bin in the histogram of Fig. 1C. The frequency corresponding to uniform usage of the different codons (1/64) is marked by the *dashed line*. There is a continuous variation of frequency/rank relation from genomes with weak (*purple lines*) to genomes with strong bias (*red lines*). We also find that there is a cascading decay of frequency with rank in genomes with strong bias.

Genomes with similar frequency/rank characteristics differ in how much they use particular different codons(Fig. 1A,B). For assessing codon usage *across* genomes, we define a non-parametric measure of ‘diversity’, *D* (eq. 4 in Methods). Diversity quantifies the frequency variation of each one of the different codons of a set of genomes with respect to the codon’s average frequency in the set. We compute diversity *D* for the genomes in each bin of the bias histogram (Fig. 1C) and plot it versus bias *B* (Fig. 2A, *gray shaded*).

**FIG. 2.**
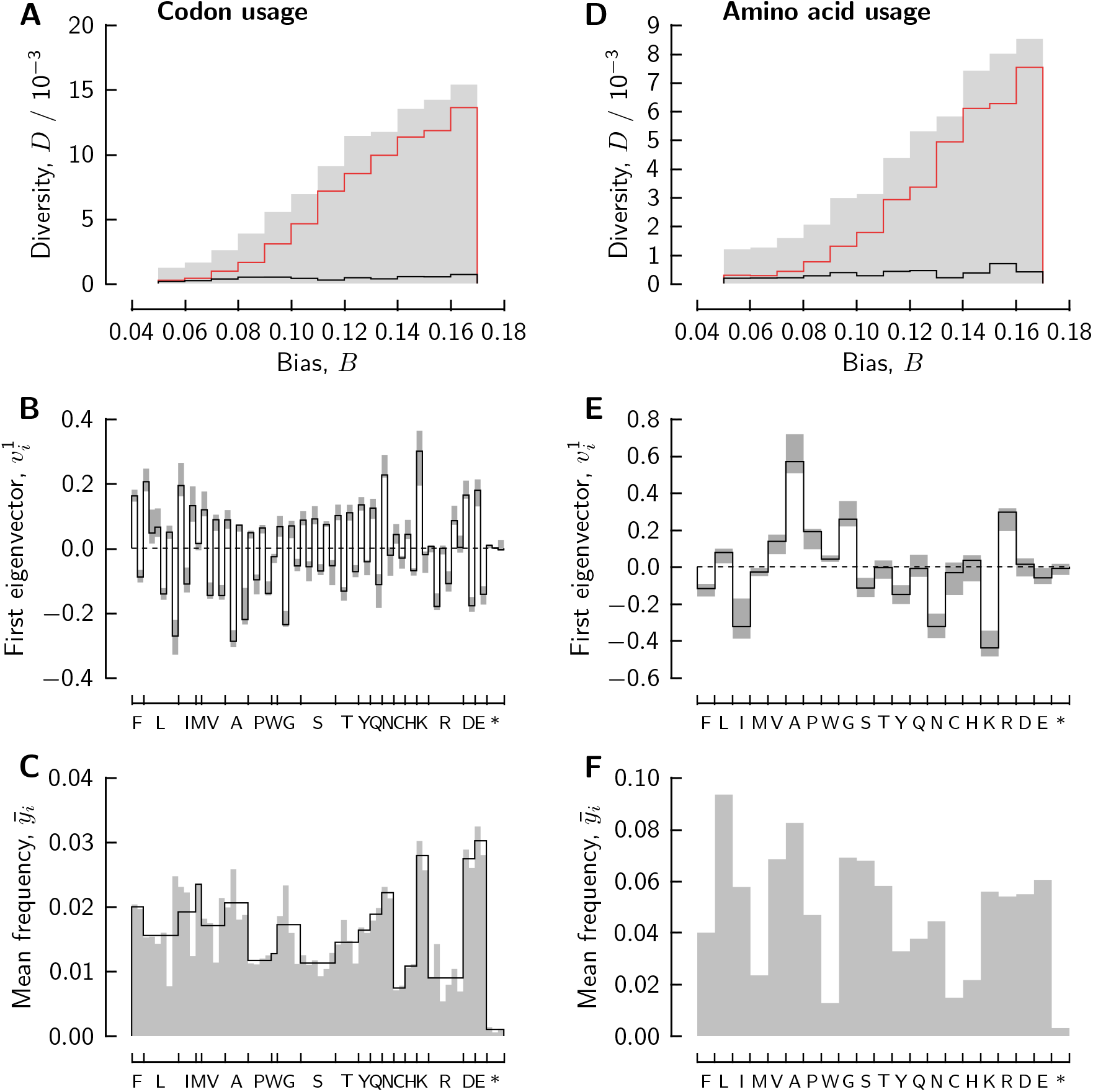
Diversity in the usage of different codons across genomes examined, by PCA. A-C: analysis of codon frequencies; D-F: analysis of amino acid frequencies. A,D: total diversity *D* computed for the genomes of each bin in Fig. 1C (*gray shade*); the first and second PCA eigenvalues are shown as red and black lines. B,E: the first PCA eigenvector components computed over the whole set of 2730 genomes (line), or over the genomes of the second to last individual bins of A,D (the gray bars indicate the range of the eigenvectors of the different bins). C,D mean frequencies computed over the full set of 2730 genomes (gray shaded); the line in C shows hypothetical mean frequencies for uniform usage of synonymous codons.

Diversity increases by about an order of magnitude with increasing bias: usage of individual different codons becomes more diverse across genomes as bias within genomes increases. That observation, already evident in the examples of Fig. 1A,B, is universal across the 2730 analyzed genomes.

The magnitude of diversity observed across genomes with strong bias B is very large. That magnitude is a strong constraint for interpretations of the diversity. Consider the set of genomes that have been grouped by their internal codon usage bias into one bin with the two genomes of Fig. 1B. These two genomes use virtually exclusive subsets of different codons. Assume for the sake of argument that one genome uses only one half of the different codons with uniform frequency (1/32), and the other genome only the other half. Assume also that one half of all genomes in the bin follow the usage pattern of one of the two genomes, and the other half of genomes follow the pattern of the other one of the two genomes. Eqs. 3 and 4 then yield the diversity *D* ≈ 1/64, which is close to the observed value found in Fig. 2A. Thus, the magnitude of the observed diversity does not only require mutually exclusive patterns of codon usage, but also that the set of genomes be divided about equally between the two alternate patterns. The variations of codon usage across the genomes of that set must involve joint variations of virtually all codon frequencies.

To investigate the extent of joint variations of different-codon frequencies across the genomes we apply principal component analysis (PCA) to the covariances of frequencies defined by eq. 5 in Methods. PCA identifies patterns of joint variations (of frequencies) and their contributions to the total variance (diversity). The linear combinations defining joint variations are computed as the unit eigenvectors of the covariance matrix, and the variance contributions as the respective eigenvalues.

The lines in Fig. 2A represent the largest and second-largest diversity components (eigenvalues) determined by PCA. Each genome set locating to one of the bins of the bias histogram (Fig. 1C) is analyzed separately, so that there are distinct eigenvalues for each individual bin. The contribution of the first principal component (*red line*) follows the total diversity (*gray shaded*) as the bias within each genome increases, such that the first principal component essentially accounts for the increase of the observed total diversity. In contrast, the second principal component (*black line*) contributes a smaller part of the total diversity that remains small as codon usage bias and total diversity increase. PCA of codon frequencies yields up to 63 non-zero eigenvalues. The total contributions of the second to the last principal components are given by the difference between the total diversity (*gray shaded*) and the contribution of the first principal component (*red line*). Because the bias-dependent part of the diversity is largely accounted for by the first principal component, we focus on that component.

Fig. 2B summarizes joint frequency variations reported in the first principal component. The coefficients of frequency variation are represented in the vertical dimension for each of the 64 different codons (the horizontal axis is divided into groups of codons that code for the indicated amino acid or stop signal (*asterisk*); codon order within the synonymous groups is listed in Table 1). A global analysis, comprising all 2730 genomes binned in Fig. 1C, yields the linear combination represented by the *solid line*. Bin for bin analyses of the genomes in the second to last bin yield combinations whose coefficients fall within the range marked by the *gray bars*.

**Table 1.** Codon frequencies† of Streptomyces griseus and Clostridium tetani

|  |  |  |  |  |  |  |  |  |  |  |  |
| --- | --- | --- | --- | --- | --- | --- | --- | --- | --- | --- | --- |
| F | UUU | 1 | 39 | W | UGG | 14 | 6 | H | CAU | 2 | 11 |
|  | UUC | 28 | 5 | G | GGU | 8 | 19 |  | CAC | 21 | 2 |
| L | UUA | 0 | 52 |  | GGC | 58 | 4 | K | AAA | 1 | 77 |
|  | UUG | 3 | 8 |  | GGA | 7 | 34 |  | AAG | 20 | 22 |
|  | CUU | 2 | 17 |  | GGG | 20 | 6 | R | CGU | 5 | 1 |
|  | CUC | 42 | 1 | S | UCU | 1 | 18 |  | CGC | 38 | 0 |
|  | CUA | 1 | 11 |  | UCC | 20 | 5 |  | CGA | 3 | 1 |
|  | CUG | 56 | 1 |  | UCA | 1 | 15 |  | CGG | 33 | 0 |
| I | AUU | 1 | 33 |  | UCG | 14 | 1 |  | AGA | 1 | 25 |
|  | AUC | 30 | 4 |  | AGU | 1 | 18 |  | AGG | 3 | 5 |
|  | AUA | 1 | 67 |  | AGC | 14 | 5 | D | GAU | 4 | 46 |
| M | AUG | 17 | 25 | T | ACU | 1 | 21 |  | GAC | 55 | 7 |
| V | GUU | 2 | 23 |  | ACC | 41 | 5 | E | GAA | 10 | 63 |
|  | GUC | 48 | 1 |  | ACA | 1 | 20 |  | GAG | 46 | 13 |
|  | GUA | 3 | 31 |  | ACG | 18 | 2 | * | UAA | 0 | 2 |
|  | GUG | 30 | 6 | Y | UAU | 1 | 35 |  | UAG | 0 | 1 |
| A | GCU | 3 | 22 |  | UAC | 20 | 6 |  | UGA | 3 | 0 |
|  | GCC | 78 | 4 | Q | CAA | 1 | 18 |  |  |  |  |
|  | GCA | 5 | 23 |  | CAG | 26 | 4 |  |  |  |  |
|  | GCG | 45 | 2 | N | AAU | 1 | 55 |  |  |  |  |
| P | CCU | 2 | 12 |  | AAC | 19 | 10 |  |  |  |  |
|  | CCC | 26 | 2 | C | UGU | 1 | 9 |  |  |  |  |
|  | CCA | 1 | 12 |  | UGC | 8 | 2 |  |  |  |  |
|  | CCG | 33 | 1 |  |  |  |  |  |  |  |  |

<sup>†</sup> Scaled by 10<sup>3</sup> and rounded to nearest integer

The first principal component thus defines variations of codon frequency that are linked among most different codons. Linkage among codon frequencies is spread much wider than that expected, e.g., from the swapping of synonymous codons for one amino acid, which is not necessarily expected to be tied to codon-swapping in the synomymous groups of other amino acids. The few different codons for which the variation is small include those encoding the single-codon amino acids (methionine, tryptophan) and the stop signal. Codons belonging to the same synonymous group typically undergo extensive, alternate variations of frequency.

While the total diversity of the genomes varies substantially with intra-genomic bias (Fig. 2A), the pattern of joint variation revealed by the first principal component is independently and consistently detected in the genomes of each of the second to the last bins of intragenomic bias. Diversity of codon usage across genomes thus is dominated by a universal pattern of variation in the codon frequencies.

The covariances studied in PCA imply that the frequency of a different codon is shifted by the average frequency of that codon in the set of genomes (eq. 5). Fig. 2C shows the average codon frequencies computed over all 2730 genomes (*gray shaded*). In many cases, the frequencies of synonymous codons have similar values (the hypothetical situation of exactly uniform usage of the synonyms is represented by the *line* in Fig. 2C to assist the eye). The variations described by the first principal component then are about the codon usage expected from the average frequencies of encoded amino acids, with quite uniform recruitment of synonymous codons.

To analyse diversity in amino acid usage that is implied in the diversity of codon usage, we translate codon frequencies of genomes into amino acid frequencies (using the standard genetic code) and submit the amino acid frequencies to PCA. The results are shown in Fig. 2D-F, side-by-side with the codon-level PCA results. Again, the first principal component describes much of the total diversity, and the associated linear combination of frequency variations is universal for the genomes of the database. We find oppositely directed changes in the frequencies of the different amino acids within some groups defined by side-chain nature. Fig. 2F shows the average amino acid frequencies about which the described variations of frequency occur. An analogous analysis, with codons grouped together by side-chain nature of the encoded amino acids, reveals a substantial shift among amino acids with non-polar versus polar side chains (Fig. S1 in Supplementary Materials).

A first principal component accounting for much of the observed diversity provides a concise means to characterize a genome *k*: the projection of observed codon frequencies onto the direction of the first principal component (eq. 6 of Methods). That projection, *z_k_*, is plotted versus genome bias B_k_ in the panels of Fig. 3. Each panel presents the genomes of the indicated CUTG database section(s). The first principal component was determined over the 2730 genomes of all panels. The genomes used as examples in Fig. 1A,B are marked by *filled red circles*. The graphs provide a perspective of genomic codon usage in the form of a map. The individual genomes of the different taxa consistently group into a V-shaped formation. That formation reveals how codon usage groups genomes into two converging lineages.

The *lines* in Fig. 3 represent the theoretical function relating codon usage bias *B* and projection *z* of codon frequencies (eq. 8). The function is derived by inserting codon frequencies *y_i_*^1^ reconstructed from eq. 7 of Methods for a given value of z into eq. 2. Observed projections *z* tend to be associated with larger values of codon usage bias *B* than those expected from the function. The difference quantifies contributions to codon usage bias that are described in the second and higher principal components. The interior of the V-formation is sparsely populated at elevated values of bias as genome locations follow the curve based on the first principal component. Thus, the description of codon usage by the first principal component becomes more accurate with increasing bias in codon usage. The variations of codon frequency described by the first eigenvector then dominate diversity.

**FIG. 3.**
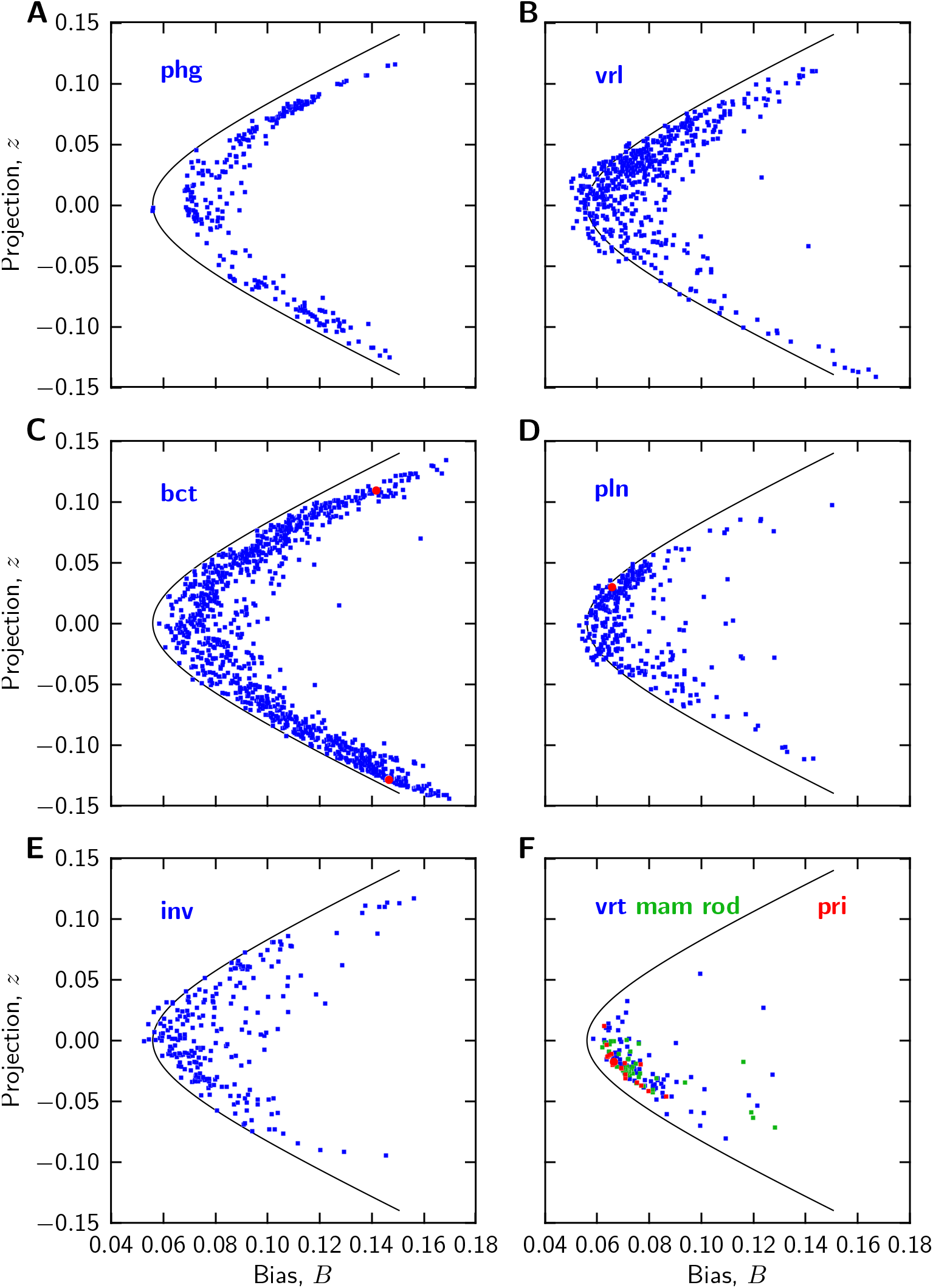
Codon usage maps of genomes. Genomes are grouped by CUTG database sections as indicated (see Methods); the section labeled ‘bct’ includes both bacterial and archaeal genomes. Red filled circles mark the genomes of *Homo sapiens* (F), *Arabidopsis thaliana* (D), *Streptomyces griseus* (C, lower circle), and *Clostridium tetani* (C, upper circle). The position of a genome relates measures of codon usage bias within the genome (horizontal axis) and of codon usage diversity with respect to average usage in the genomes (vertical axis). The curves are computed from eq. 8 of Methods and discussed in the text. The Supplementary Materials include a separate file for each CUTG database, ‘xxx_map.txt’, tabulating species identifier, species name, and map coordinates *B* and *z*.

Fig. 4A-D shows examples of individual codon frequencies that have been reconstructed from the first principal component (*lines,* based on eq. 7 of Methods). These are superimposed on the observed frequencies (*gray shaded*) of the four genomes introduced as examples in Fig. 1A,B. The comparison suggests that the partial PCA information used for the reconstructions captures the diverse codon usages of these genomes quite well, in particular those of the two bacterial genomes. The human and plant genomes involve relatively small bias values *B_k_* and first-axis projections *z_k_* (Fig. 3D,F). Hence, their reconstruction is chiefly determined by the average codon usage *ȳ_i_* (Fig. 2C), whereas the reconstructions of the two bacterial genomes strongly depend on the variations described by the first principal component.

**FIG. 4.**
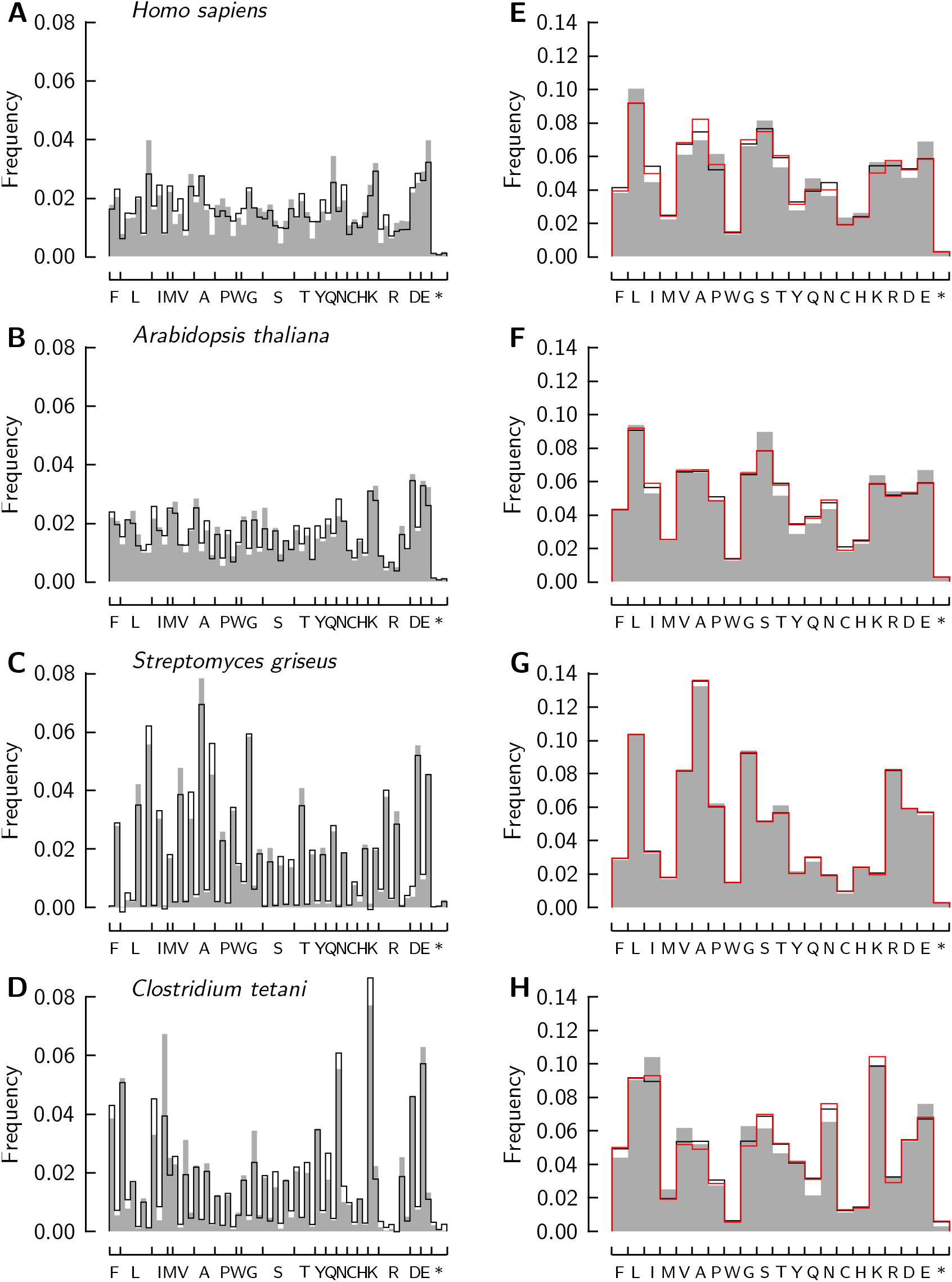
Codon (A-D) and amino acid frequencies (E-H) reconstructed from PCA results. Gray shaded: observed codon frequencies; black lines: frequencies reconstructed from the first principal component. Example genomes: *Homo sapiens* (A, E), *Arabidopsis thaliana* (B, F), *Streptomyces griseus* (C, G), and *Clostridium tetani* (D, H). The red lines in E-H are amino acid frequencies computed from the reconstructed codon frequencies of A-D. All reconstructions are made with the PCA results obtained from the genomes in the bias bin to which the example genome locates (see Figs. 1C and 3).

An analogous reconstruction is made for the amino acid frequencies using the PCA results shown in Fig. 2C-E. Fig. 4E-G show the amino acid frequencies derived from the observed codon frequencies (*gray shaded*), the frequencies reconstructed from the PCA of the amino acid frequencies (*black lines*), and the amino acid frequencies calculated from the reconstructed codon frequencies (*red lines*). The amino acid frequencies reconstructed from the first principal components of the two sets of PCA results are closely similar - the variation in amino acid frequencies is also captured in the PCA results of the individual codon frequencies. Hence, the vari-ations among genomes that are observed in the usage of synonymous codons and the variations that are observed in the usage of particular amino acids are tightly linked.

## DISCUSSION

The main result of our study of 2730 genomes from different taxa can be summarized as follows: Codon usage varies both within and between genomes to different degrees but a universal pattern links the frequency variations of the different codons across genomes. That pattern is revealed by the first principal component of the frequency covariance matrix. The codon usage of indvidual genomes can be mapped in two coordinates, one a non-parametric measure of bias of codon usage within a genome, the other the weight of the first principal component for the genome. In that map, genomes group together in a V-shaped formation, with the majority of genomes locating near the junction of the formation (at small bias and small weight), and fewer genomes with weights diverging in two branches toward larger bias (Fig. 3).

To identify the pattern of codon usage we now consider two specific genomes whose codon frequencies are juxtaposed for comparison in Table 1. *Streptomyces griseus* and *Clostridium tetani E88* are recognized in the map as being among the organisms with the most biased and diverse codon usages (Fig. 3C). Indeed, *Streptomyces* and *Clostridium* use nearly exclusive subsets of the 64 diferent codons (Fig. 1B). Inspection of Table 1 reveals that the two bacteria have strong preferences for different synonymous codons. That preference concerns the third base: *Clostridium* prefers synonyms with U or A as third base, *Streptomyces* prefers synonyms with G or C as third base. This way, each bacterium restricts its codon usage virtually to one half of the genetic code table.

Codon usage generally defines two lineages of genomes, one preferring U/A as third base of codons (positive branch in Fig. 3), the other preferring G/C (negative branch in Fig. 3). Genomes locating to the regions of maximal spread between the branches in essence restrict their codon usage to either half of the genetic code table, whereas the genomes locating to the junction of the branches approach recruitment of the full code table. The vertical dimension of the map thereby approximately represents the ‘UA3 content’ of a genome, which is the complement of the GC_3_ content often quoted in the litrature. Fig. S2 in Supplementary Materials uses UA_3_ content itself as the vertical dimension of genome maps, and shows that maps essentially identical to those in Fig. 3 are directly constructed from the codon frequencies of the genome set.

A second kind of bias in codon usage is observed in Table 1 in the encoding of the amino acids leucine (L) and arginine (R). Each of these amino acids is represented by six synonymous codons, using two different patterns in the first two bases. *Streptomyces* uses codons with one pattern in the first two bases, whereas *Clostridium* uses codons with the other pattern (Table 1). These two kinds of variation in synonymous codon usage are accompanied by a substantial variation in amino acid usage (Figs. 4G,H).

Together, the three different aspects that render the codon and amino acid usages in *Streptomyces* and *Clostridium* distinct from each other represent a universal pattern of diversity. That pattern dominates the variation of codon and amino acid usage in the entire set of genomes that we have sampled. The pattern is found from the genomes with strong bias in their codon usage to the genomes that approach uniform codon usage (Fig. 2B). It involves proportionate, joint variations of the frequencies of most different codons.

A mutational hypothesis for the origin of codon usage bias has been widely discussed [2–4]. Codon usage studied in 100 eubacterial and archaeal genomes using PCA has been interpreted to be “constrained by genomewide mutational processes” [17]. Since the pattern of frequency variation that we find in the first principal component of a larger sample of organisms does not show dispersion as the extents of bias and diversity vary, a theory based on point mutations would require mutation rates of different codons to be in fixed ratios to one another while varying in absolute magnitude among different groups of organisms. A theory based on point mutations of unconstrained rates is unconstrained in the crucial parameters that would determine the codon frequencies to be explained.

Variations of codon usage that are described by a linear combination (eq. 7, Figs. 3,4) are expected from a mixing, and re-mixing, of gene populations that make two distinct usages of bias and/or amino acids. We therefore propose lateral DNA transfer [18, 19]
as the main mechanism for the variations of codon usage among organisms. Lateral gene acquisitions of large scale have recently been found to occur between the eubacterial and archaeal domains [20].

Lateral DNA transfer would naturally account for other phenomena as well. Transferred genes would carry with them preferences for the third base of codons if they cross the lineages of genomes apparent in Fig. 3. GC_3_ content is known to be diverse across the genes within individual genomes (e.g., [21, 22]). Lateral DNA transfer between organisms with diverse codon and amino acid usages is expected to level genome-wide codon usage bias and to balance genome-wide usage of amino acids. Lateral DNA transfer, thus, is consistent with convergence of codon-usage lineages over time (Fig. 3).

Genomes with strong bias of codon usage are found among the archaea and eubacteria that share the largest numbers of homologous proteins attributed to the hypothetical last universal common ancestor [8, 9]. Such genomes populate the most divergent parts of our codon usage map (Fig. 3C; panel C represents both eubacteria and archaea as these are not distinguished as separate sections in the CUTG database; see file ‘bct_map.txt’ in Supplementary Materials for the included individual genomes). For instance, *Methylobacterium extorquens* (Archaea) and *Halobacterium salinarum* (Archaea) share preference for using G/C as third base with *Anaeromyxobacter dehalogenans* (Protobacteria), *Actinoplanes fruilensis* (Actinobacteria), and *Desulfovibrio vulgaris* (Delta-protobacteria). On the other hand, *Methanosphaera stadtmanae* (Euryarchaeota) and, as described, *Clostridium tetani* (Clostridia) prefer U/A as third bases. While these organisms share homologous proteins that follow vertical but not lateral lineages of primary structure [9], they form two genetic lineages that encode proteins with distinct subsets of the genetic code table. Two genetic lineages among organisms thought to be close to ancestral forms of life might complicate phylogenetic inferences. On the other hand, codon usage and amino acid composition themselves might provide information complementary to amino acid sequences in phylogenetic studies.

A divergence of the encoding of amino acids might have evolved in organisms over the billions of years after their putative common ancestors lived. It is, however, difficult to imagine evolutionary pressures that might have produced two antipodal usages of the genetic code, with a neat separation between the usages of U and A, or G and C, as third bases of codons. (Neat separation is, e.g., not necessary for reducing the required number of different tRNAs by one half). If the phenomenon, however, is ancestral, it makes the unknown origin of the genetic code even more mysterious. Two histories need to be unraveled: that of a universal code for amino acids, and that of a dichotomy in the usage of redundancy.

## CONCLUSIONS

The variation of codon usage across genomes is primarily a genome-wide linked variation between the usages of either U and A, or G and C, as the third bases of codons. The variation among different synonymous codons is accompanied by a variation among different amino acids. These variations across genomes are like those expected from lateral DNA transfer among organisms. The variation of codon usage divides organisms into two lineages that span different taxa. The two genetic lineages are most diverse for prokaryotes sharing homologous proteins ascribed to the hypothetical last universal common ancestor (LUCA), but converge toward uniform codon usage for multicellular organisms.

---

**FIG. S1.**
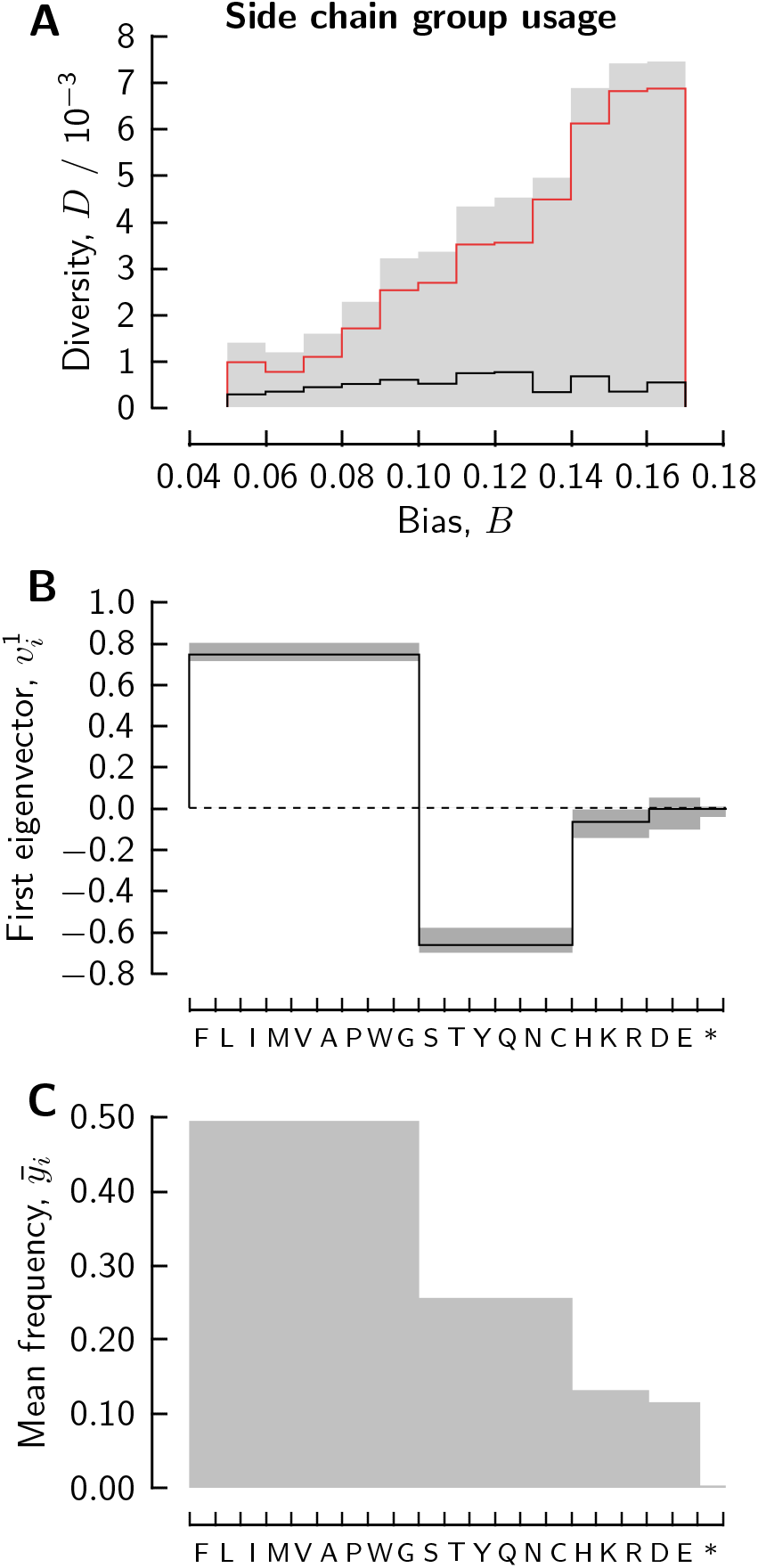
Usage of different amino acids grouped by chemical properties. Group frequencies were computed by summing the frequencies of codons for all amino acids of the groups (nonpolar, polar, basic, acidic, and stop signal) and subjected to PCA. **A**: total diversity *D* computed for the genomes of each bin in Fig. 1C (*gray shade*); the first and second PCA eigenvalues are shown as red and black lines. **B**: the first PCA eigenvector components computed over the whole set of 2730 genomes (line),or over the genomes of the second to last individual bins of panel A (the gray bars indicate the range of the eigenvectors of the different bins). **C**: mean frequencies computed over the full set of 2730 genomes (gray shaded). See Fig. 2 of text for the corresponding analysis of codon and (individual) amino acid frequencies.

**FIG. S2.**
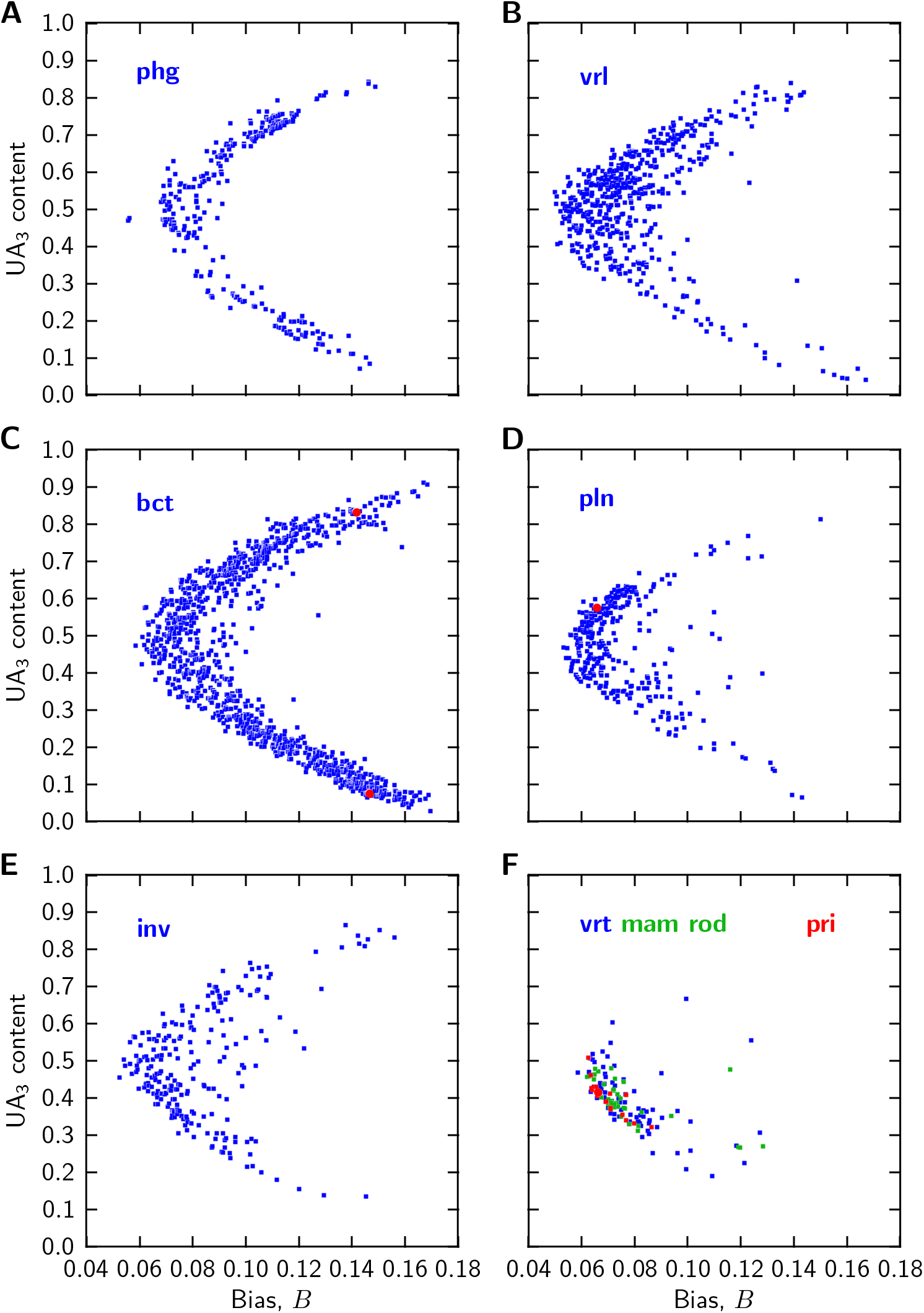
Codon usage maps of genomes. Genomes are grouped by CUTG database sections as indicated (see Methods); the section labeled ‘bct’ includes both bacterial and archaeal genomes. Red filled circles mark the genomes of *Homo sapiens* (F),*Arabidopsis thaliana* (D), *Streptomyces griseus* (C, lower circle), and *Clostridium tetani* (C, upper circle). The position of a genome relates a measure of codon usage bias within the genome (horizontal dimension) and UA_3_ content (vertical dimension), where UA_3_ content is the complement of GC_3_ content. See Fig. 3 of text for the corresponding maps based on PCA. UA_3_ content is also tabulated (as last column) in the Supplementary Materials, files ‘xxx_map.txt’.

